# The alleviation of neuropathic pain behaviours by a single injection of a synthetic substance P-botulinum conjugate persists for up to 120d and can be restored with a second injection of conjugate

**DOI:** 10.1101/2020.12.28.423185

**Authors:** Maria Maiarù, Charlotte Leese, Bazbek Davletov, Stephen P. Hunt

**Affiliations:** Department of Pharmacology, School of Pharmacy, University of Reading, Room 104, Hopkins Building, Whiteknights Campus, Reading, United Kingdom, RG6 6UB; Department of Biomedical Science, Firth Court, University of Sheffield, Sheffield S10 2TN; Cell and Developmental Biology, Medawar Building, University College London, Gower Street, London, WC1E 6BT, UK

## Abstract

There is an urgent need for new pain-relieving therapies. We have previously shown using mouse models of persistent pain that a single intrathecal injection of substance P conjugated to the light chain of botulinum toxin (SP-BOT) silenced neurons in the dorsal horn of the spinal cord and alleviated mechanical hypersensitivity. The SP-BOT construct selectively silenced neurokinin 1 receptor positive (NK1R+) neurons in the superficial dorsal horn of the spinal cord. A subset of these NK1R+ neurons are nociceptive projection neurons and convey injury-related information to the brainstem, initiating and maintaining programmes of escape and recovery essential for healing. Previously, we observed a reduction in mechanical hypersensitivity in a spared nerve injury (SNI) model of neuropathic pain state after intrathecal injection of SP-BOT over the lumbar spinal cord and lasting for up to 40 days. In this latest study, we have extended these observations and now show that thermal and affective measures of pain behaviour were also alleviated by a single intrathecal injection of SP-BOT. By introducing SNI 30 days, 60 days, 90 days or 120 days after injection of SP-BOT we have established that NK1R+ spinal neurons in the superficial lamina of the dorsal horn were silenced for up to 120 days following a single intrathecal injection of the botulinum construct. We also show that behavioural alleviation of neuropathic pain symptoms could be reinstated by a second injection of SP-BOT at 120 days. Taken together this research demonstrates that this recently developed botulinum toxin conjugate provides a powerful new way of providing long term pain relief without toxicity following a single injection and also has a therapeutic potential for repeated dosing when pain begins to return.

## Introduction

Noxious stimulation of sufficient intensity to induce tissue damage leads to increased excitability of peripheral and central neuronal circuits that heightens pain experience and serves to protect damaged tissue from further trauma[27; 45; 57]. In some cases, on-going disease or the failure of potentiated pain signaling networks to return to pre-injury levels leads to chronic pain conditions[34; 58]. Chronic or persistent pain is highly prevalent and extremely difficult to treat with prescribed drugs such as opioids often having significant unwanted side effects. Although research into developing new analgesic drug therapies has been intense, translating knowledge from preclinical observations in animal models to new therapies in the clinic has been challenging[24].

Previous research has identified neuronal networks within the dorsal horn and pathways from the dorsal horn to the brain as important for the regulation of persistent pain states[25]. More recently the contribution of ascending pathways from the superficial dorsal horn of the spinal cord to the brain have also been shown to be essential for the maintenance of persistent pain states[13; 31; 33; 38; 42; 43; 47]. Many of these superficial spinal projection neurons express the substance-P (SP) preferring neurokinin 1 receptor (NK1R)[50] and can be ablated with intrathecal (i.t.) application of substance P-saporin (SP-SAP) conjugates[38; 42] resulting in normal acute thermal and mechanical stimulus detection but a loss of mechanical and thermal hypersensitivity in inflammatory and neuropathic pain models. SP-SAP gains specific access to NK1R-positive neurons through binding of SP to the G-protein coupled NK1 receptor, triggering internalization of the receptor and the transport of bound SP-SAP complex into the cell. Unfortunately, while SP-SAP treatment was shown to alleviate bone cancer pain in companion dogs[7], clinicians were cautious about taking the approach forward to the clinic primarily because the toxin conjugate killed neurons and clinical grade constructs proved difficult to synthesize. We therefore designed alternative constructs using botulinum neurotoxin serotoxin A (BoNT/A) which silenced but did not damage neurons[4; 23; 36; 37]. BoNT/A is made up of a light-chain zinc endopeptidase and a heavy chain that is responsible for binding the toxin to neuronal receptors and promoting essential light-chain translocation across the endosomal membrane. Once internalized within the neuron, the light chain has the capacity to silence neurons via the specific proteolytic cleavage of synaptosomal-associated protein 25 (SNAP25), a protein essential for synaptic release[49]. SNAP25 is found in neurons but not in glial cells[29] and is the unique substrate for botulinum protease cleavage. By exploiting a recently introduced protein ‘stapling’ method’ we linked the light-chain/translocation domain (LcTd) of botulinum neurotoxin type A (BOT) to neurotransmitter ligand SP to target key pain-processing neurons in the dorsal horn[36].

With this botulinum-based approach[36] a single intrathecal injection of the neuropeptide substance P conjugate (SP-BOT) was shown to alleviate inflammatory and neuropathic pain states through the silencing of nociceptive pathways for up to 40 days. Importantly, the intrathecal injection of SP-BOT construct had no effect on baseline mechanical thresholds in naïve mice. Furthermore, using NK1R gene knockout mice we showed that the NK1R was essential for SP-BOT–mediated reduction of mechanical hypersensitivity in the spared-nerve injury (SNI) model of neuropathic pain. Using immunohistochemistry to detect the cleaved SNAP25 (cSNAP25) we found that SP-BOT was internalized only by NK1R-expressing neurons. In addition, no changes in the extent of NK1R+ immunofluorescence were found in the dorsal horn of mice that had been treated with SP-BOT, suggesting lack of construct-induced cytotoxicity or receptor down-regulation. We also utilized c-Fos histochemistry to confirm a loss of activity in the ascending pathway to the brain in SP-BOT treated and stimulated mice directly implicating spinal to parabrachial NK1R+ projection neurons as targets for the construct[36].

The present study was designed to examine the length of time over which a single injection of SP-BOT was effective in reducing the symptoms of neuropathic pain by investigating mechanical and cold thermal responses and affective motivational behaviours. We found that neuropathic pain behaviours were alleviated for up to 120 days after a single intrathecal injection of SP-BOT and importantly that analgesia could then be reinstated by a second injection of the conjugate.

## Results

### Pre-emptive injection of SP-BOT prevents the full development of neuropathic pain state

To establish the long-term efficacy of the SP-BOT conjugate we adopted an approach that minimizes unnecessary discomfort for the mouse by assessing the impact of a neuropathic pain model generated at 30, 60, 90 and 120 days *after* intrathecal injection of botulinum-neuropeptide conjugates (Fig. 1). Throughout our recent studies intrathecal injections of botulinum constructs were generally given *after* the inflammatory or neuropathic pain state had been established, usually 3 days after, and the behaviour measured for 14-28 days. Here we made intrathecal injections of SP-BOT in naïve mice *before* establishing a neuropathic pain state (Fig 1).

**Fig 1:**
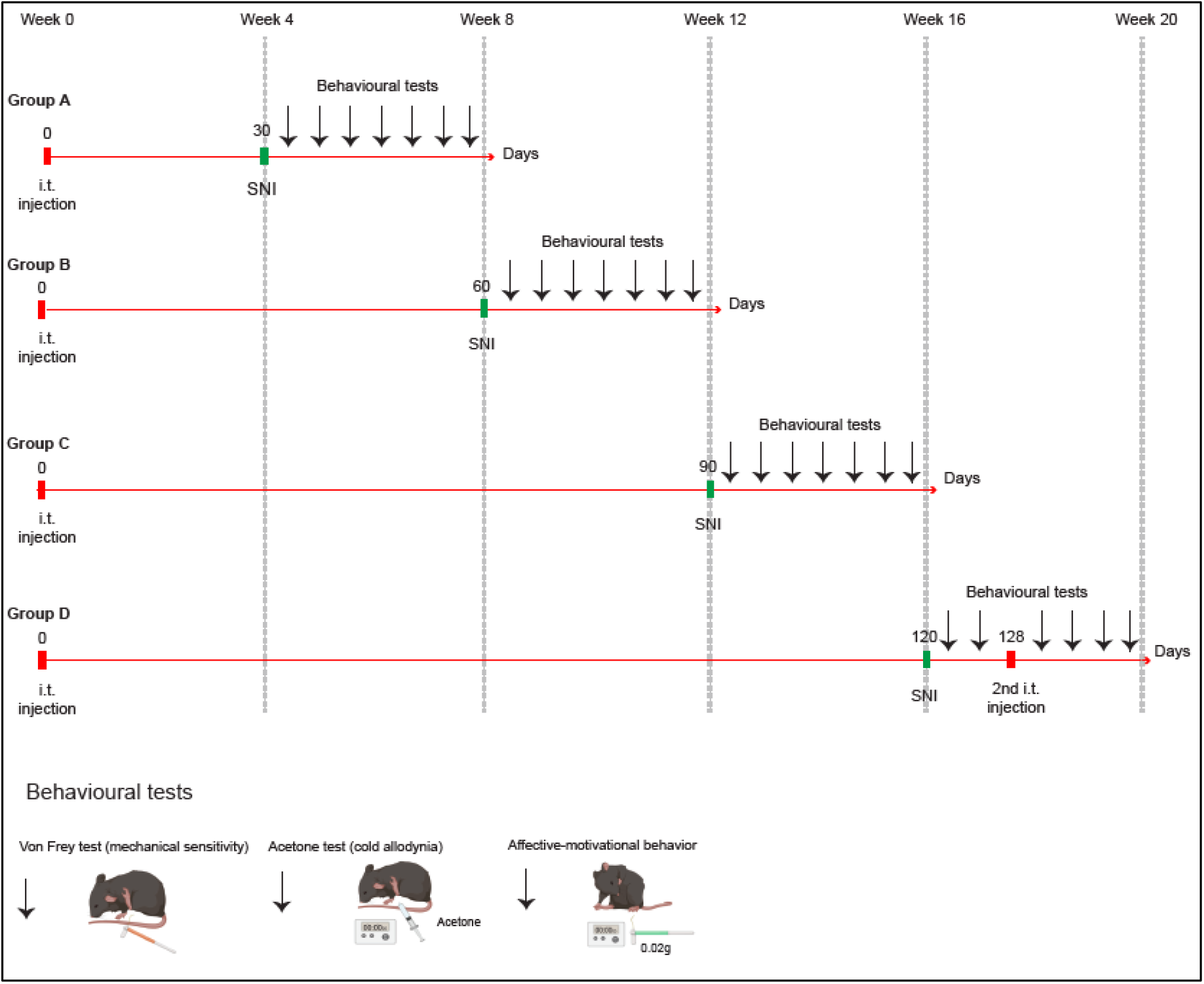
Plan of experiment. Four groups of mice (A-D) were subject to spared nerve injury (SNI) surgery (green marker) at 30 (Group A), 60 (Group B), 90 (Group C) or 120 days after intrathecal injection of SP-BOT (100 ng/3 μl) or saline vehicle (3 μl) at day 0. Group D received a 2^nd^ intrathecal injection 8 days after SNI surgery (128 days after 1^st^ intrathecal injection). Behavioural testing for mechanical sensitivity, cold allodynia and affective motivational behaviour was carried out before and after SNI (black arrows). Mechanical thresholds were assessed using calibrated von Frey filaments, cold allodynia was measured using the acetone drop test and affective-motivational behaviour was assessed by the cumulative response to a one-second von Frey stimulation (0.02g) of the hind paw over a 30s period.

Mechanical and cold allodynia are common symptoms of neuropathic pain. In patients with neuropathic pain a light touch can induce severe pain that is difficult to treat using conventional drugs, such as NSAIDs or opioids [24; 26; 32; 55]. Here, we used a model of neuropathic pain induced by nerve injury, the SNI model, and monitored responses to a number of different stimuli. Using Von Frey filaments, we showed that a pre-emptive injection of SP-BOT 30, 60 and 90 days before the induction of the pain state was able to prevent the full development of mechanical allodynia observed after nerve injury (Fig 2, A-B-C). Furthermore, using a commonly used model to test cold allodynia in mice, the acetone evaporation test, we showed a reduction in the licking/biting behaviour in mice injected with SP-BOT 30, 60 and 90 days before the injury, compared with saline-injected mice (Fig 3, A-B-C).

**Fig 2:**
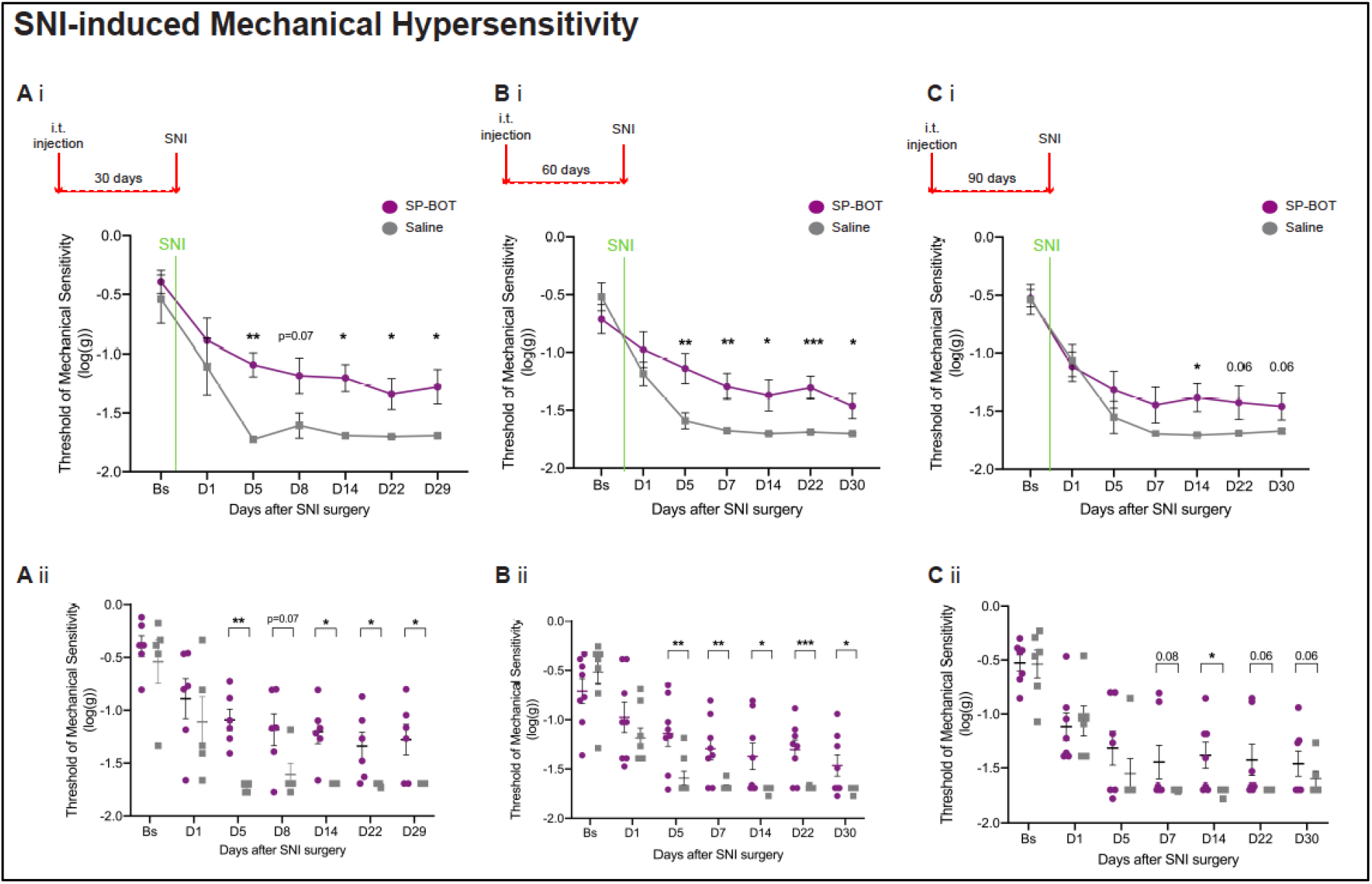
Pre-emptive injection of SP-BOT is effective for more than 100 days in reducing mechanical sensitivity that develops after nerve injury. **A (i-ii).** Mice in Group A received i.t. injection of SP-BOT (100ng/3 μl) or saline 30 days before SNI surgery and were then tested for up to 29 days after surgery. Mechanical threshold assessed using calibrated von Frey filament in mice before (Bs, Baseline) and after SNI surgery (D1 to D29). Two-way ANOVA, factor TREATMENT day 1 (D1) to D29: F _1,9_=10.12; P=0.011. n=6/5. **B (i-ii).** Mice in Group B received i.t. injection of SP-BOT or saline 60 days before SNI surgery and were then tested for up to 30 days after surgery. Mechanical threshold assessed using calibrated von Frey filament in mice before (Bs) and after SNI surgery (D1 to D30). Two-way ANOVA, factor TREATMENT day 1 (D1) to D30: F _1,14_=15.43; P=0.002. n=8/8. **C (i-ii).** Mice in Group C received i.t. injection of SP-BOT or saline 90 days before SNI surgery and were then tested for up to 30 days after surgery. Mechanical threshold assessed using calibrated von Frey filament in mice before (Bs) and after SNI surgery (D1 to D30). Two-way ANOVA, factor TREATMENT day 5 (D5) to D30: F _1,11_=4.91; P=0.049. n=7/6. Data show means ± SEM. *P < 0.05, **P < 0.01, ***P < 0.001.

**Fig 3:**
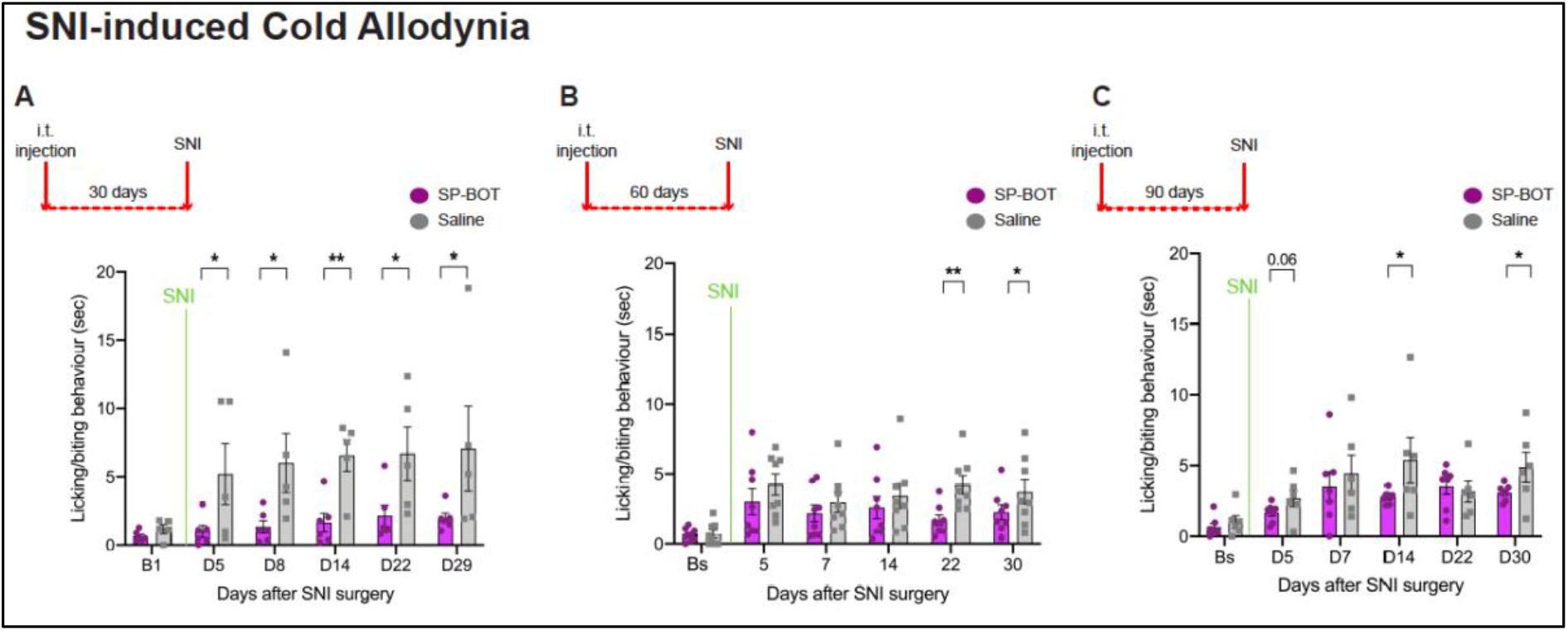
Pre-emptive injection of SP-BOT is effective for up to 120d in reducing the cold allodynia that develops after nerve injury. **A (i-ii).** Mice in Group A received i.t. injection of SP-BOT or saline 30 days before SNI surgery and were then tested for up to 29 days after surgery. Cold allodynia assessed using acetone drop applied to the hindpaw ipsilateral to the injury (left) in mice before (Bs) and after SNI surgery. Two-way ANOVA, factor TREATMENT day 5 (D5) to D29: F _1,9_=19.45; P=0.002. n=6/5. **B (i-ii).** Mice in Group B received i.t. injection of SP-BOT or saline 60 days before SNI surgery and were then tested for cold allodynia for up to 30 days after surgery. Two-way ANOVA, factor TREATMENT day 14 (D14) to D30: F _1,14_=5.23; P=0.038. n=8/8. **C (i-ii).** Mice in Group C received i.t. injection of SP-BOT or saline 90 days before SNI surgery and were then tested for cold allodynia for up to 30 days after surgery. n=7/6. Data show means ± SEM. *P < 0.05, **P < 0.01.

Pain is a complex experience, including not only nociceptive and nocifensive but also affective-motivational components[14; 15; 58]. We measured the affective-motivational component of nociceptive behaviour by monitoring the responses to a single von Frey filament over 30sec which unlike acute reflex responses to stimulation requires participation of forebrain circuits[15; 59]. Pre-emptive intrathecal treatment with SP-BOT did not change affective-motivational responses when measured 30, 60 and 90 days after injection (Fig 4, A-B-C, (Bs) Baseline measure). By contrast, induction of the pain state produced an increased affective-motivational response in saline-injected mice but a significantly weaker response in SP-BOT injected mice (Fig 4, A-B-C).

**Fig 4:**
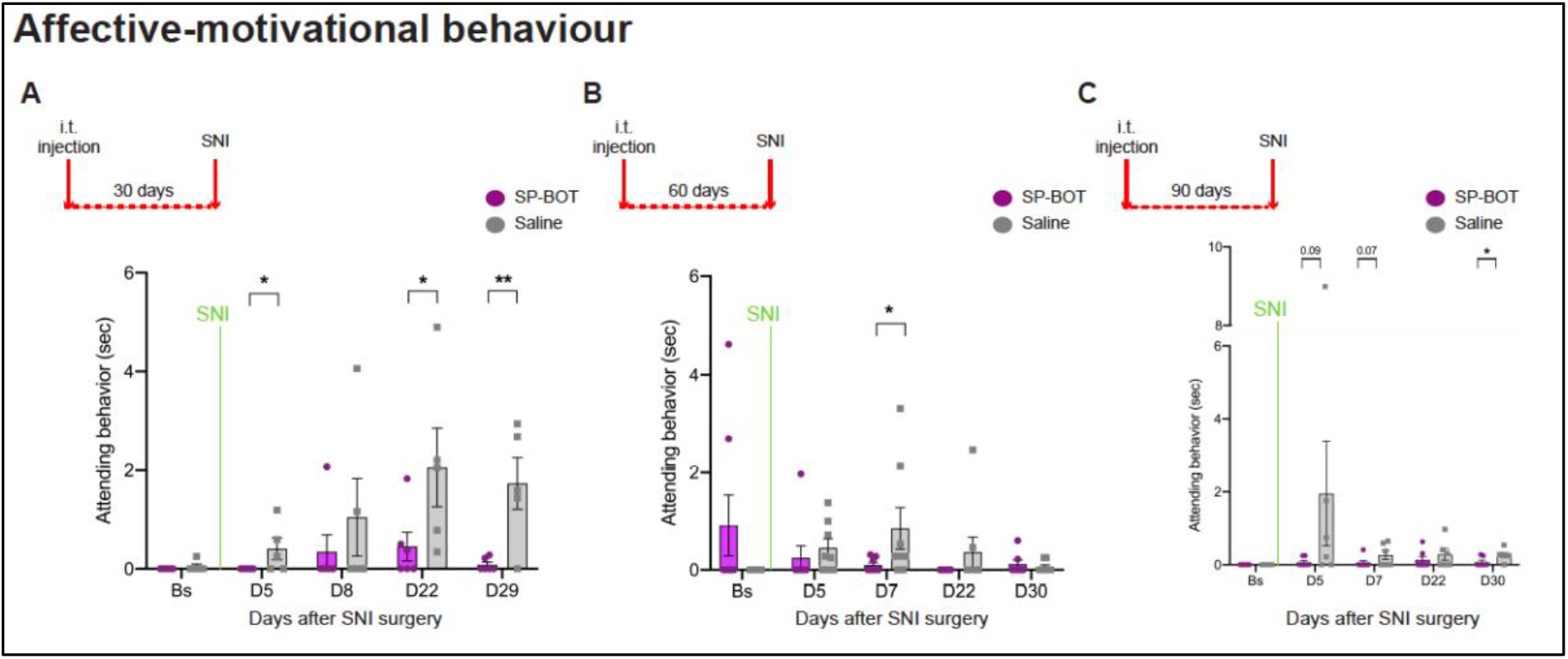
Pre-emptive injection of SP-BOT is effective for around 120d in attenuating the development of mechanically-induced affective-motivational behaviours that develop after nerve injury. Time course of affective-motivational behaviour in response to a one-second von Frey stimulation (0.02g) before (Bs) and after SNI surgery. **A.** Effect of pre-emptive injection of SP-BOT and saline vehicle 30 days before surgery. Two-Way ANOVA, factor TREATMENT day 5 (D5) to D29: F_1,9_=12.64; P=0.006. n=6/5. **B** Effect of pre-emptive injection of SP-BOT and saline vehicle 60 days before surgery. Two-Way ANOVA, factor TREATMENT day 5 (D5) to D22: F_1.14_=4.49; P=0.05. n=8/8. **C.** Effect of pre-emptive injection of SP-BOT and saline vehicle 90 days before surgery. Two-Way ANOVA, factor TREATMENT day 7 (D7) to D30: F_1.11_=6.48; P=0.027. n=7/6. Data show means ± SEM. Overlaid points are individual subject scores. *P < 0.05, **P < 0.01.

Interesting, we observed a decline in responsiveness in the saline group when the pain state was induced a longer time point (60 and 90 days after i.t. injection). This reduction was observed both in the affective-motivational behaviour and in cold allodynia scores, suggesting that a longer period in the animal house after the i.t. injection or age[41] may contribute to the development of pain behaviour. However, significant differences in pain scores between SP-BOT and saline-injected mice were still present.

### SP-BOT-induced analgesia can be restored by a second intrathecal injection

A pre-emptive injection of SP-BOT 120 days before SNI was no longer able to reduce the symptoms of the neuropathic pain state (Fig 5). However, a second injection of SP-BOT 8 days after SNI surgery was able to reduce mechanical hypersensitivity and the affective components of pain behaviour (Fig 5 B). Moreover, after SNI surgery we observed a further development of cold allodynia compared to baseline measure in saline-injected mice but not in SP-BOT injected mice (Fig 5 C), implying that the construct was able to alleviate all components of the pain state investigated (Fig 5 D).

**Fig 5:**
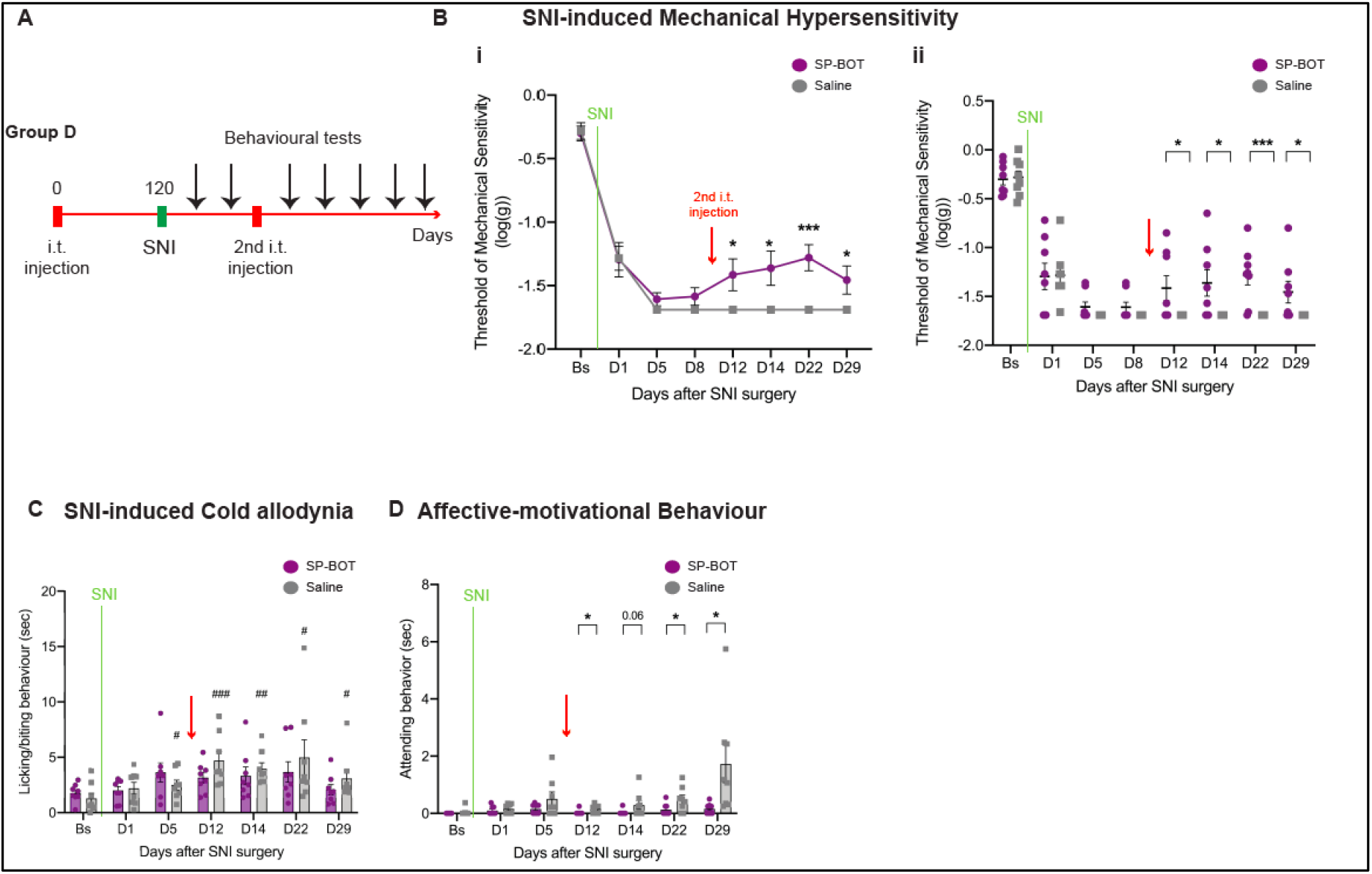
The analgesic impact of the first SP-BOT injection on neuropathic pain was lost by 120d but restored by a second intrathecal injection of SP-BOT. **A.** Mice in Group D (see Fig. 1) received i.t. injection of SP-BOT or saline 120 days before SNI. Behavioural tests were performed for up to 8 days after surgery following which mice received a second i.t. injection of SP-BOT or saline vehicle. Mechanosensitivity, cold allodynia and affective motivational behaviours were measured for up to 30 days after SNI surgery and reinjection. (Black arrows). **B.** (i-ii) Mechanical thresholds assessed in mice at baseline (Bs) and after SNI surgery were similar in both treated and control groups. The second injection of SP-BOT however resulted in reduced mechanical hypersensitivity in the SP-BOT group. Two-way ANOVA, factor TREATMENT day 1(D1) to D29: F1,13=5.86; P=0.031. Two-way ANOVA, factor TREATMENT D12 to D29: F _1,14_=12.07; P=0.004. n=8/8. **C.** Cold allodynia assessed in mice before (Bs) and after SNI surgery and following the second injection of SP-BOT. Cold allodynia developed in saline-injected mice but not in SP-BOT injected mice. Student’s t test was performed to compare saline group after SNI to baseline. Two-way ANOVA, factor TREATMENT D12 to D29: F1,14=2.47; P=0.14. n=8/8. **D.** Affective-motivational behaviours before (Bs) and after SNI surgery were similar but attenuated in mice that received a second injection of SP-BOT. Two-way ANOVA, factor TREATMENT D12 to D29: F _1,14_=11.72; P=0.004. n=8/8. Data show means ± SEM. Overlaid points are individual subject scores. N=8/8, *P<0.05, **P<0.01 (Student’s t test). * SP-BOT vs Saline; # Saline BS vs Saline-post surgery.

We also found that pre-emptive treatment with SP-BOT was as effective as injections of the construct made after establishing the pain state. Pre-emptive SP-BOT had a maximal possible effect (MPE) on reducing SNI-induced mechanical hypersensitivity of 28.2%. In the present experiment when the injection of SP-BOT followed SNI the MPE was 28% or 24.8% in a previously published experiment[36] (Fig 6).

**Fig 6:**
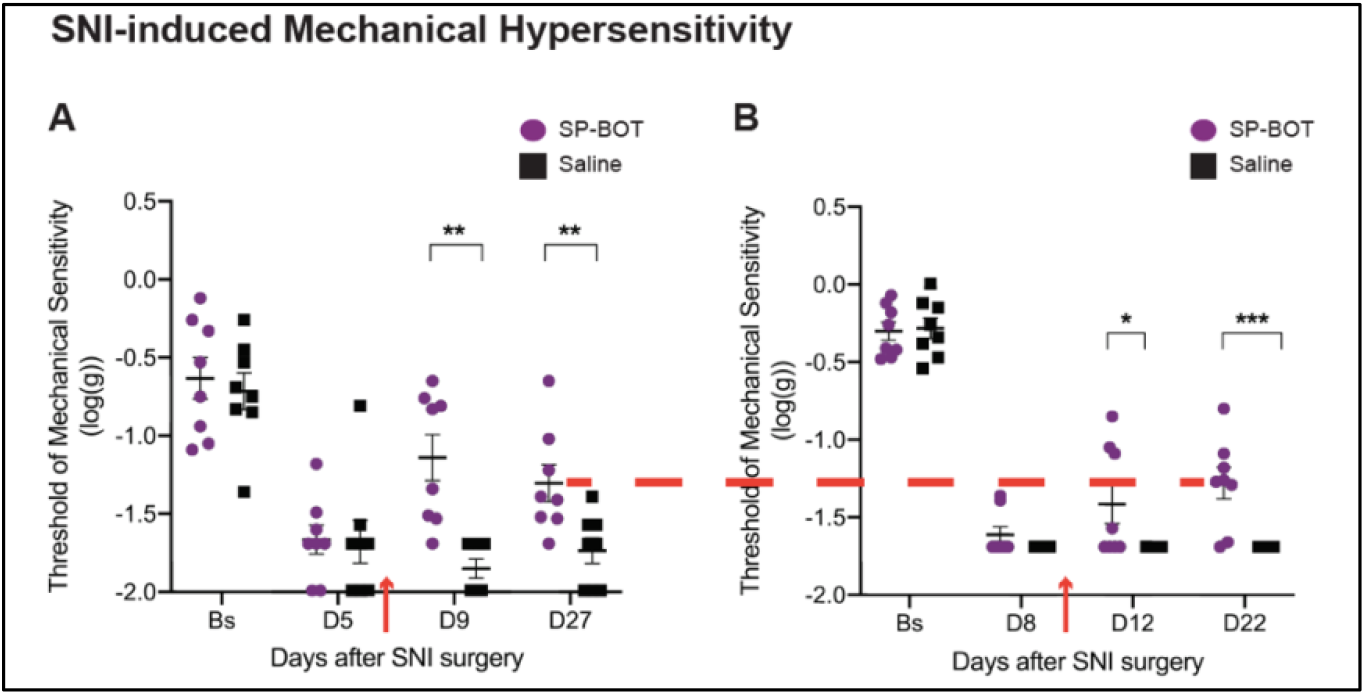
Reinjection of SP-BOT induces the same analgesic effect on neuropathic pain as in our previous study. **A.** Data from our previous study (Maiarú et al, 2018). Mice received i.t. injection of SP-BOT (100ng/3 ml) (red arrow) or saline 5 days after SNI. N=8/8. **B**. Data from this study. Mice received i.t. injection of SP-BOT (100ng/3 ml) (red arrow) or saline 8 days after SNI. N=8/8. Data show means ± SEM. Overlaid points are individual subject scores. *P<0.05, **P<0.01, ***P<0.001 (Student’s t test). * SP-BOT vs Saline.

## Discussion

We previously demonstrated that novel substance P-botulinum constructs could alleviate chronic neuropathic and inflammatory pain states in mice following a single intrathecal injection of the conjugate. Specificity was demonstrated by showing that these conjugates were only internalised by NK1R expressing neurons and were ineffective in NK1R knockout mice[36]. Here we determined the duration of silencing of NK1R+ neurons by introducing SNI at various time points *after* i.t. injection of SP-BOT. Using an extended range of behavioural assays including thermal and affective motivational measures of pain behaviour we found that the botulinum conjugates reduced pain scores for over 100 days. We also demonstrated that a reduction in pain scores could be re-established by a second injection of SP-BOT at 128 days after the first injection.

The time over which neuronal silencing could be observed was comparable to that previously reported in rats using botulinum toxin A (BoNT/A) injections into the hind limb[6; 17]. Once internalized within the motor neuron, the light chain of BoNT/A silenced peripheral motor neurons for several months via the specific proteolytic cleavage of SNAP25, a protein essential for synaptic release. This inhibition was slowly reversed as the endopeptidase lost activity and function was usually restored in rats within 100 days. Data on direct injections of BoNT/A into the central nervous system are more limited but recent studies have shown that the functional consequences of injections of BoNT/A directly into the striatum of hemi-Parkinsonian rats were also reversed within 90-100 days but residual activity was seen for much longer, up to one year[3]. No toxicity has been reported following BoNT/A injections into peripheral tissues including motor neurons and clinically the process is frequently repeated. We show here that a second injection of SP-BOT, re-administered once the effect of the first injection on pain behaviour has subsided, alleviates chronic neuropathic pain and to the same extent as shown in our previous reports when SP-BOT was given *after* SNI. This also implies that the NK1R+ neurons, that are the substrate for silencing and the alleviation of pain states, are present and have not been lost or remodeled during the initial lengthy period of silencing. In contrast, motor neuron recovery from BoNT/A silencing is accompanied by the temporary appearance of functional nerve sprouts that facilitated restoration of function[17].

The function of lamina I NK1R+ neurons has come under increasing scrutiny. The majority of lamina I projection neurons terminate in the parabrachial area and spino-parabrachial neurons (SPNs) account for 5% of lamina I neurons in the rat lumbar enlargement[8; 28; 50; 52]. 90% of these cells were NK1R+, and with generally larger cell bodies than other NK1R+ neurons in this lamina[2; 11; 19; 50; 51]. However up to 45% of lamina I neurons express the NK1R and the majority of these lamina 1 NK1R+ neurons are thought to be excitatory interneurons[51]. The great majority of spinoparabrachial projection neurons (SPN) are nociceptive specific, and almost all are activated by noxious mechanical, chemical or thermal stimuli[5; 13; 20; 21; 39; 53; 54]. Both interneurons and projection neurons receive convergent cutaneous and visceral afferent input[35]. The situation appears to be similar for both rat and mouse[8; 28]. Evidence also suggests that a small number of SPNs survive SP-saporin (SP-SAP) injection[48]. These may be the populations of non-NK1R SPNs including so called giant neurons[44] and those identified by their expression of the orphan receptor GPR83[13]. These SPNs may be functionally separate and mediate distinct behaviours[13].

Recent studies have claimed that parabrachial neurons and, by implication SPNs, are essential for many of the behaviours associated with injury, including escape behaviours, analgesia and the formation of aversive memories as well as affective-motivational behaviours and extensive grooming associated with anxiety [12; 13; 30; 56]. However, the licking and guarding behaviours that following noxious heat stimulation were unaffected by ablation of NK1+ neurons with SP-SAP but the operant escape response was substantially reduced[56]. Licking and biting responses also occur in decerebrate rats[59] implying that these were responses to noxious stimulation requiring only intact brainstem but not forebrain function. But how necessary SPNs are for ‘reflex responses’ that occur immediately after injury is unclear. SP-SAP ablation studies have shown that NK1R+ SPNs are essential for the full expression of first and second phases of capsaicin-induced nocifensive behaviours but only for the second phase of formalin induced behaviours[38; 42].

However, these findings arise largely from studies of acute pain. In chronic pain states mechanical and thermal sensitivity are reduced by ablation or silencing of superficial NK1R+ neurons[42; 47; 48; 56]. Thus, Mantyh and colleagues demonstrated that the NK1R+ SPNs were essential for the full expression of chronic pain behaviours caused by nerve damage or inflammation[38; 42]. Ablation of these neurons with SP-SAP conjugates alleviated pain states when given either before or after establishing spinal nerve ligation and indeed has been shown to be an effective treatment in alleviating pain in companion dogs with bone cancer [7; 42]. SP-SAP ablation also reduced chronic itch and cold allodynia seen after spinal nerve ligation[1; 9; 47; 48]. It has been concluded that increased excitability of spinal nociceptive networks following injury is maintained by spinal-brainstem loops. Thus, the evidence suggests that the ascending NK1R+ pathway initially triggers escape responses to noxious stimulation and continues to inform the brain about the extent and severity of injury. In turn the brain modulates spinal excitability through descending brainstem-spinal pathways [21; 33; 47; 48]. Silencing of lamina 1 NK1R neurons with SP-BOT replicates the impact of SP-SAP on the alleviation of chronic pain states by restricting the flow of information concerning injury to the brain.

It is unclear if SP-BOT also silences the large population of lightly labelled small diameter NK1R+ interneurons found in lamina I as well as the smaller group of larger PBNs[8; 51]. Labelling with fluorescent-SP conjugates failed to reveal more than a few labelled cell bodies within the superficial spinal cord[53; 54] and no substantial cell death or gliosis was detected in lamina I of SP-SAP treated rats[38; 42; 47; 56]. Taken together this suggests that SP conjugates preferentially target the larger NK1R+ expressing neurons of lamina I that are for the most part SPNs[11; 46].

Our study builds on a previous research implying that NK1R+ projection neurons that project to the brainstem signal the existence of bodily injury and initiate and manage the programmes of escape, recovery and recuperation essential for healing[57; 58]. Evidence also suggests that NK1+ spinal parabrachial neurons maintain chronic pain states notably in the presence of an unresolved injury such as nerve damage. The clinical problem of managing chronic pain whether it be from on-going disease, nerve damage or from previous injury is immense and has proved extremely challenging to deal with. We show here that a single intrathecal injection of a SP-BOT conjugate provides long term pain relief by silencing a key pain-signalling pathway to the brain and with the added advantages that the procedure is repeatable and non-toxic to cells of the spinal cord. The translatable appeal of this approach is undeniable and this new approach may provide substantial alleviation from chronic pain states in both patients and animals.

## Methods

### Mice

Subjects in all experiments were adult male mice, C57BL/6, (8 to 12 weeks old) from Charles River (UK). All mice were kept in their home cage in a temperature-controlled (20° ± 1°C) environment, with a light-dark cycle of 12 hours (lights on at 7:30 a.m.). Food and water were provided *ad libitum*. All efforts were made to minimize animal suffering and to reduce the number of animal used (UK Animal Act, 1986).

### Design and purification of botulinum constructs

Each BoNT/A consists of three domains: the binding domain, the translocation domain, and the catalytic light-chain domain, a zinc metallopeptidase. We used a protein stapling technique to produce light chain-translocation domain (LcTd) conjugated to substance P. The synthesis that has been described previously for SP with in vitro controls for specificity is detailed in (Arsenault et al, 2013). Briefly, to synthesize the constructs, first, fusion protein consisting of the LcTd of the botulinum type A1 strain was fused to SNAP25 (LcTd-S25) and was prepared as previously described [16; 23]. The chemically synthesized syntaxin-SP peptide had the sequence Ac-EIIKLENSIRELHDMFMDMAMLVESQGEMIDRIEYNVEHAVDYVE-Ahx-Ahx-RPKPQQFFGLMNH2, where Ahx stands for aminohexanoic acid. Second, the protein “staple” was prepared recombinantly from the rat vesicle-associated membrane protein 2 (VAMP2) sequence (amino acids 3 to 84) inserted into the XhoI site of pGEX-KG. Oriented attachment of peptides to protein was achieved by the SNARE assembly reaction. LcTd-S25, VAMP2 (3 to 84), and syntaxin-SP were mixed at a molar ratio of 1:1:1 in 100 mM NaCl, 20 mM Hepes, and 0.4% n-octylglucoside at pH 7.4 (buffer A). The assembly mic was left at 20°C for 1 hour to allow formation of the irreversible SNARE ternary complex. SDS-resistant and irreversibly assembled protein conjugates were visualized using Novex NuPAGE 12% bis-tris SDS–PAGE (polyacrylamide gel electrophoresis) gels (Invitrogen) run at 4°C in a NuPAGE MES SDS running buffer (Invitrogen). All recombinant proteins were expressed in the BL21-Gold (DE3) pLysS strain of Escherichia coli (Agilent) in pGEX-KG vectors as glutathione S-transferase (GST) C-terminal fusion proteins cleavable by thrombin. GST fusion constructs were purified by glutathione affinity chromatography and cleaved by thrombin. Synthetic peptides were made by Peptide Synthetics Ltd. All constructs were tested in neuronally-differentiated SiMa neuroblastoma cell cultures and cleavage of SNAP25 determined using Western Blotting as previously described[36].

### Intrathecal injections

Intrathecal injections were performed in naïve mice under isoflurane anaesthesia[22]. Mice were held firmly but gently by the pelvic girdle using thumb and forefinger of the nondominant hand. The skin above the iliac crest was pulled tautly to create a horizontal plane where the needle was inserted. Using the other hand, the experimenter traced the spinal column of the mouse, rounding or curving the column slightly to open the spaces between vertebrae. A 30-gauge needle connected to a 10-μl Hamilton syringe was used to enter between the vertebrae. After injection, the syringe was rotated and removed, and posture and locomotion were checked as the animal regained consciousness. All intrathecally delivered drugs and vehicle solutions were injected in a 3-μl volume.

### Pain model

#### Mouse neuropathic model: Spared Nerve Injury (SNI)

At different time points after the i.t. injection (see Fig. 1), mice underwent SNI surgery. The SNI was performed as previously described[18]. Briefly, under isoflurane anaesthesia, the skin on the lateral surface of the thigh was incised, and a section made directly though the biceps femoris muscle exposing the sciatic nerve and its three terminal branches: the sural, the common peroneal, and the tibial nerves. The common peroneal and the tibial nerves were tight-ligated with 5-0 silk suture and sectioned distal to the ligation. Great care was taken to avoid any contact with the spared sural nerve. Complete hemostasis was confirmed, and the wound was sutured.

### Behavioural testing

Three behavioural assays were used to detect changes in nociceptive behaviours: von Frey filaments to measure acute withdrawal from the stimulus, affective-motivational responses to a single von Frey that involved directed licking and biting of the paw and lifting or guarding of the paw. Paw withdrawal reflexes are thought to involve spinal cord and brainstem circuits while affective-motivational responses are complex behaviours thought to indicate motivation and arousal. Finally, we assessed cold allodynia a hall mark of neuropathic pain. The experimenter was always blind to treatment group for all behavioural tests.

### von Frey filament test (Mechanical Sensitivity)

Mice were placed in Plexiglas chambers, located on an elevated wire grid, and allowed to habituate for at least 1 hour. After this time, the plantar surface of the paw was stimulated with a series of calibrated von Frey’s monofilaments. The threshold was determined by using the up-down-method [10]. The data are expressed as log of the mean of the 50% pain threshold ± SEM. Maximal Possible Effect (MPE) was calculated as previously described [36].

### Affective-Motivational behaviour

The protocol described by Corder and colleagues[15] was adapted to evaluate mechanical-induced affective-motivational response. Mice were placed in Plexiglas chambers, located on an elevated wire grid, and allowed to habituate for at least 1 hour. A 0.02g von Frey filament was applied for 1 second to the hindpaw, and the duration of attending behaviour (direct licking and biting of the paw, extended lifting or guarding of the paw) was collected for up to 30 seconds after the stimulation.

### Acetone test (Cold Allodynia)

For assessment of cold sensitivity, the acetone test was used as previously described[18]. After habituation in Plexiglas chambers, located on an elevated wire grid, a drop (50 μl) of acetone was applied to the plantar area of the hind paw ipsilateral to the site of injury, avoiding mechanical stimulation of the paw with the syringe. Total time licking/biting of the hindpaw was recorded with an arbitrary maximum cut-off time of 20 seconds.

### Data and Statistical analysis

All experiments were randomized and performed by a blinded researcher. All statistical tests were performed using the IBM SPSS Statistic programme (version 20), and P < 0.05 was considered statistically significant. For the behavioural experiments, statistical analysis was performed on the data normalized by log transformation (von Frey data), as suggested by Mills *et al* [40]. Difference in sensitivity was assessed using repeated measures two-way or one-way ANOVA, as appropriate and as indicated. In all cases, “time” was treated as within-subjects factor and “treatment” was treated as between-subject factor.

The MPE was calculated as:

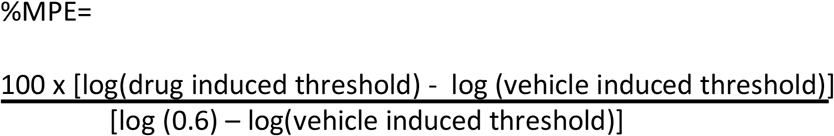

where log(0.6 g) is our maximum von Frey’s force applied.

Please note that, as in our previous study [36], we logged the data of von Frey filaments test to ensure a normal distribution because the von Frey’s hairs are distributed on an exponential scale. Mills *et al* [40] recently demonstrated that log transformation makes more “mathematical and biological sense”.

## Acknowledgments

We thank Sandrine M Geranton, Sara Hestehave, Sofia Fontana-Giusti and Silvia Silva-Hucha for comments on the manuscript.

## Author contributions

M.M., B.D., and S.P.H. designed experiments. C.L., and B.D. designed and synthesized botulinum constructs. M.M. conducted all behavioural experiments. M.M. and S.P.H. analyzed data. M.M. and S.P.H. wrote the manuscript.

## Data and materials availability

All data associated with this study are present in the paper

